# Information-theory-based benchmarking and feature selection algorithm improve cell type annotation and reproducibility of single cell RNA-seq data analysis pipelines

**DOI:** 10.1101/2020.11.02.365510

**Authors:** Ziyou Ren, Martin Gerlach, Hanyu Shi, GR Scott Budinger, Luís A. Nunes Amaral

## Abstract

Single cell RNA sequencing (scRNA-seq) data are now routinely generated in experimental practice because of their promise to enable the quantitative study of biological processes at the single cell level. However, cell type and cell state annotations remain an important computational challenge in analyzing scRNA-seq data. Here, we report on the development of a benchmark dataset where reference annotations are generated independently from transcriptomic measurements. We used this benchmark to systematically investigate the impact on labelling accuracy of different approaches to feature selection, of different clustering algorithms, and of different sets of parameter values. We show that an approach grounded on information theory can provide a general, reliable, and accurate process for discarding uninformative features and to optimize cluster resolution in single cell RNA-seq data analysis.

## Introduction

The application of single cell RNA sequencing (scRNA-seq) as a hypothesis generation step is becoming increasingly popular in biology and medical research because of the promise to (i) efficiently identify novel cell types from transcriptomic data (Patel et al. 2014; Villani et al. 2017); (ii) identify heterogeneity within defined cell populations during homeostasis and disease (Reyfman et al. 2019; Lawson et al. 2015); (iii) follow cellular lineage trajectories(Treutlein et al. 2014; Rizvi et al. 2017; Collin et al. 2019); or (iv) identify novel genetic markers for disease (Miyamoto et al. 2015; Min et al. 2015; Lindström et al. 2019).

A key step in scRNA-seq analysis is cell type identification using computational clustering algorithms. While supervised clustering approaches, i.e. identifying cell types based on reference dataset, have been developed to use the data produced by large international efforts such as the Human Primary Cell Atlas (HPCA) and the Encyclopedia of DNA Elements (ENCODE) (Aran et al. 2019; Mabbott et al. 2013; Davis et al. 2018), many studies still rely on unsupervised clustering approaches i.e. identifying cell types based on expression patterns. Among all clustering algorithms, Louvain method (Blondel et al. 2008) is the most popular method in scRNA-seq because of its high efficiency in analyzing high dimensional data. It is the default clustering algorithm in one widely used analysis package, Seurat (Satija et al. 2015). Besides Louvain methods, these integrated packages implement a number of normalizations, feature selection and other cell clustering algorithms. Nevertheless, all unsupervised clustering algorithms try to quantify the similarity of every pairwise cell expression profile in order to identify “clusters” of cells of the same type. Clusters are then assigned to known cell types via the manual identification of genes expressed by cells in the cluster based on *a priori* biological knowledge. Visual inspection is then used to “validate” the quality of the clustering, or the clustering produced by a new algorithm is compared to the clustering reported using a previously published algorithm.

As single cell data are increasingly used to define differences in cell state, such as might develop during disease, in addition to cell type, trustworthy quantitative tools are necessary to assess whether the partitioning of cell populations by an algorithm is justified. An issue with many currents approaches is that they have not been validating against label assignment that are independent of the transcriptomic data used by the clustering algorithm. We focus here first the development of a benchmarking data set that does not rely on transcriptomic data for generating cell annotations. Specifically, we take surface protein measurements and use a decision tree model to generate cell type labels. We use the normalized mutual information to quantify the agreement between pairs of cell clustering partitions and systemically evaluate the impact of sparsity and genes on the accuracy and reproducibility of current clustering algorithms against the surface-protein-based reference labels.

Additionally, we validate an entropy based, non-parametric feature selection algorithm to evaluate the information content for genes in scRNA-seq data. Using only the genes with high information content, both the clustering accuracy and reproducibility were improved for several clustering algorithms.

## Results

To avoid circularity in comparing the performance of algorithms for clustering single cell RNA-seq data, we must generate a reference dataset that does not rely on annotations generated making use of the same RNA-seq data (Fig. 1A). A publicly available peripheral blood mononuclear cells (PBMC) dataset compiled by 10x Genomics enables us to accomplish this goal. This dataset provides protein staining with TotalSeq-B antibodies for every cell and single cell transcriptomic profiling using the 10x platform. The protein staining data allow us to assign cell type labels following the approaches used traditionally for cell classification in immunohistochemistry and flow cytometry and independently from the RNA-seq data (Fig. 1B). This benchmarking data set then enables it to systematically evaluate the impact of different feature selection algorithms and different clustering algorithms on cell labelling accuracy.

**Figure 1.**
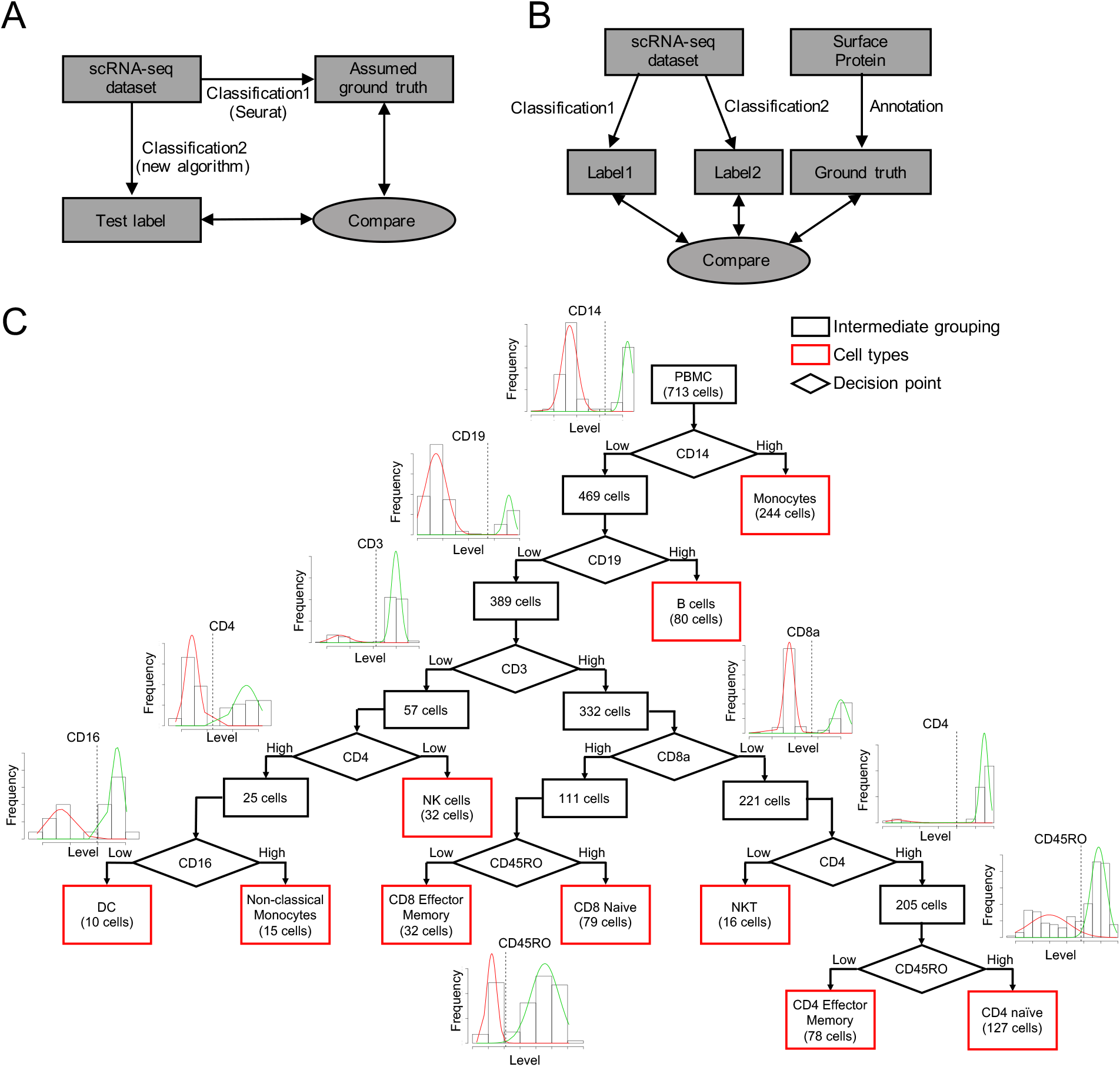
Limitations of current framework for optimizing scRNA-seq cell type classification algorithms and development of an externally validated dataset. A. Current implicit frameworks for optimizing scRNA-seq classification algorithms assume that some algorithm, typically Louvain method from Seurat, yields a ground-truth classification against which the accuracy of other algorithms is then determined. B. A reproducible, objective framework would make use of an independently-obtained, robust, and reproducible independently-generated reference annotations against which the accuracy of scRNA-seq classification algorithms can be objectively determined. We believe that in order to avoid circularity reference annotations should be based on independent approaches such as surface protein expression or immunostaining. C. Creation of externally validated labelling for peripheral blood mononuclear cells (PBMCs) from a healthy donor released by 10x Genomics (https://support.10xgenomics.com/single-cell-gene-expression/datasets/3.0.0/pbmc_1k_protein_v3). The dataset includes simultaneous single cell levels for 17 surface proteins measured using ToptalSeq-B antibodies, as well as transcriptomic profiles for 713 cells. Surface protein level from the cells is consistent with a bimodal distribution. We model empirical distribution as a mixture of two Gaussian peaks and detect a threshold for binary classification of protein level, which can be used for classification with a biologically-grounded decision tree enabling us to classify every cell into one of ten cell types.

Following the traditional approach in cell classification using surface protein markers with reproducible workflow, we produce a decision tree for cell type annotation (Fig. 1C). It is visually apparent that the distribution of the surface protein levels is consistent in all cases with a bimodal distribution. Therefore, we model the distribution using a Gaussian mixture model with two peaks and determine a binary cutoff by identifying the value for which the likelihoods of a cell belonging to each of the two groups are identical. We list the estimated parameters for the decision tree in Supplementary Table 1. We use this approach to classify every one of the 713 cells in the dataset into one of ten cell types. These cell types cover all of the well-known cell types in PBMC (Supplementary Table 2).

In order to test the robustness of our classification and to acknowledge uncertainty in a cell’s classification, we also consider the case in which we set cells with protein levels that yield odds ratios smaller than 5:1 between estimated Gaussian functions for two peaks as “unclassified cells”. Using this procedure, we are still able to classify 696 of 713 cells (Fig. S1 and Supplementary Table 3). Importantly, our results and conclusions do not change when considering this alternative labelling approach.

Next, we investigate whether our benchmarking approach enables us to quantify the impacts of data sparsity and potentially uninformative genes on the performance of clustering algorithms. For concreteness, we focus first on the performance of build-in Louvain methods in Seurat, which we will simply refer to as Seurat in the remainder of the manuscript. Seurat is one of the most widely adopted tool in scRNA-seq analysis. To test the hypothesis that high sparsity and overwhelming numbers of uninformative genes will induce poor classification accuracy, we generate synthetic datasets that allow us to vary data sparsity and the number of potentially uninformative genes.

As a first step, we apply Seurat to the surface protein level dataset in which sparsity is negligible and all features are by construction meaningful. Using the full data for the seventeen cell-surface proteins, we are able to classify cell types with very high precision (Fig. 2A-B and Fig. S2). Indeed, the only misclassifications occur in separating cells types that are relatively infrequent in the dataset. Thus, as one would expect, Seurat performs very well when applied to data with negligible data sparsity.

**Figure 2.**
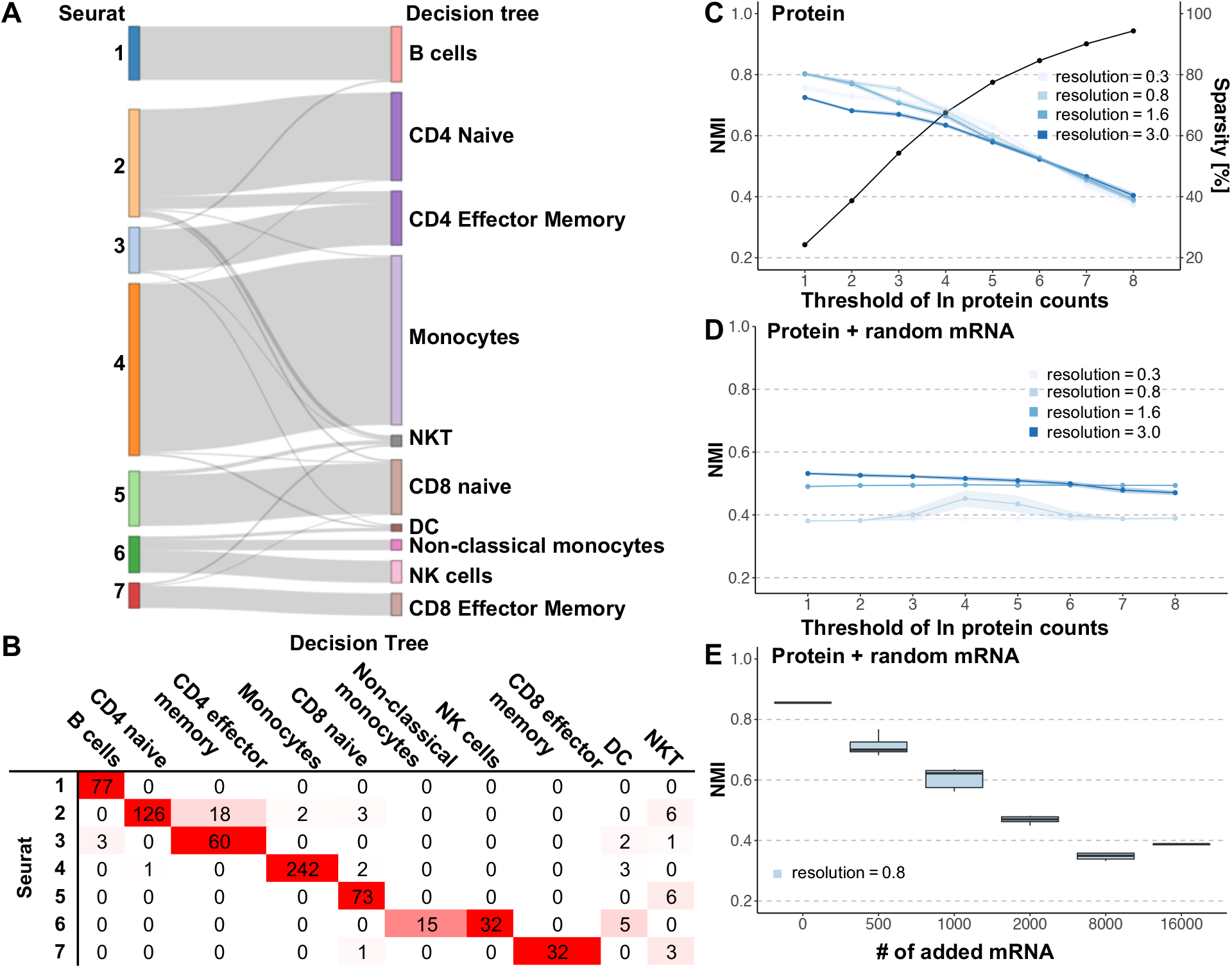
Clustering accuracy of Seurat decreases dramatically with increasing levels of data sparsity and increasing number of uninformative variables. A. Sankey diagram of Seurat classification based solely on surface protein labels. The left side shows the clusters identified by Seurat with the parameter set to the default value (0.8). The right side shows the cell types we identify using surface protein data and a decision tree. B. Confusion matrix between Seurat classification and surface protein label in A. The most likely cell type is highlighted in red on the diagonal line. C. Effect of data sparsity on accuracy of Seurat algorithm for four values of the parameter, *resolution*, ranging from 0.3 to 3.0. We model data sparsity by setting a threshold level for the detectability of the log protein counts (see text for details). As we increase data sparsity, by moving the mid-point of sigmoidal function, we find a steady decrease in accuracy. Notably, this decrease does not depend on the value of *resolution*. Sparsity measured by percentage of zero counts (black line) with the increasing threshold of log protein counts. About 90% of the counts become zero for *t* ≥8. D. Effect of number of possibly uninformative variables and data sparsity on accuracy. We repeat the procedure described above for surface protein levels, but now include in the data the measured levels of 16,000 randomly selected mRNAs. The large number of uninformative variables included in the transcriptomic data decreases clustering accuracy by nearly 40% for low levels of data sparsity. Interestingly, it yields best accuracy with *resolution* >1.6, not with the default value. E. Effect of number of possibly uninformative variables on accuracy. We generate 10 replicates by adding to the protein-levels data randomly-selected sets with data from varying numbers of mRNAs in order to estimate the confidence intervals for accuracy of classification. If none of the possibly uninformative mRNA is included, then classification accuracy measured is very high. As we include data from an increasing number of mRNAs, we find a steady decrease in accuracy across all parameter settings.

To systemically investigate how data sparsity affects clustering accuracy without the complications due to uninformative features, we model data sparsity by setting a model for the probability that a specific surface protein’s level is too low to be detected (refer to Methods for details). Compared with an available method for producing sparsity where counts are drawn from the negative binomial distribution (Zappia, Phipson, and Oshlack 2017), or a naïve model where counts are randomly changed to 0, our method better captures the physical process occurring in real experiments where the presence of a value of zero counts will be correlated with the expression level.

We systematically change the value of the threshold level *t*, and quantify the clustering accuracy using the normalized mutual information (NMI) between the independently-generated reference labels and the labels generated by the clustering algorithm on the synthetic dataset. NMI is a commonly used metric to quantify the overlap between different partition. An NMI equal to 1 indicates perfect overlap, whereas a value of 0 indicates random overlap (Witten and Frank 2002). Figure 2C shows that our benchmarking approach correctly reflects that steady decrease in clustering accuracy as data sparsity increases, that is, as the detectability threshold increases to higher levels.

Next, we examine how a large number of potentially uninformative features interacts with data sparsity to affect the accuracy of clustering algorithm as another important part of benchmarking. To this end, we add the data from 16,000 randomly selected genes from the transcriptomics dataset to the synthetic data for protein levels with distinct detectability threshold generated in the previous step (Fig. 2D). As expected, the large number of likely uninformative features leads to a decrease in clustering accuracy. Surprisingly, this decrease is even more dramatic than one would expect as it completely overshadows the impact of data sparsity.

In order to quantify the specific effect of uninformative features, we add to the full surface protein level data different numbers of randomly selected genes from the transcriptomic dataset. It is remarkable that transcriptomic data for even as few as 1,000 randomly selected genes reduces the algorithm’s NMI by 25% (from 0.8 to 0.6, Fig. 2E). This decrease clearly demonstrates that more data are not always helpful. In fact, an increase in the number of uninformative features dramatically decreases performance.

These results demonstrate that the inclusion of features that are uninformative for the purpose of cell labeling may have an unexpected impact on clustering accuracy (Kiselev, Andrews, and Hemberg 2019). Before proceeding, we must emphasize that important genes for cell specific functions are not necessarily informative of cell type if they are expressed in multiple cell populations. For example, MALAT1 and HLA-A, genes which are highly conserved or constitutively expressed, are critical for cell function (Hutchinson et al. 2007; Kovats et al. 1990), however, their broad expression across cell populations means they are uninformative for cellular classification. Due to the importance of the degree to which a gene may provide information for cell classification, we pursue an approach grounded on information theory that we have recently demonstrated can dramatically improve the performance of algorithms for the classification of texts (Gerlach, Shi, and Amaral 2019). Unlike other currently pursued approaches, such as filtering out genes with low coefficient of variation, ours has a solid theoretical grounding and does not require any assumptions about the distribution of counts of unique molecular identifiers (UMIs).

Concisely, conditional entropy *H*(*g*|*S*) is calculated for each gene given the UMI frequency. We also created null models by random distribution to calculate average conditional entropy over different realizations, 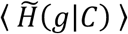 (details refer to method). The information content *I*(*g*) is defined as the difference between *H*(*g*|*S*) and 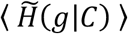. Higher *I*(*g*) indicates that the gene has high expression in a subset of cells, which will be useful to determine cell types.

To test the robustness of our filtering approach, we calculate *H*(*g*|*S*), 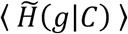, and *I*(*g*) for three PBMC datasets publicly available on the 10x Genomics website. As predicted, we observe a very strong dependency of 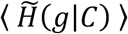 on gene total expression level (Fig. 3A). We also find that most genes have *I*(*g*) ≈ 0, indicating that those genes cannot possibly provide meaningful information concerning cell identities. Surprisingly, 15 out of 17 of the genes coding the surface protein markers we use in generating labels are not informative for determining cell type.

**Figure 3.**
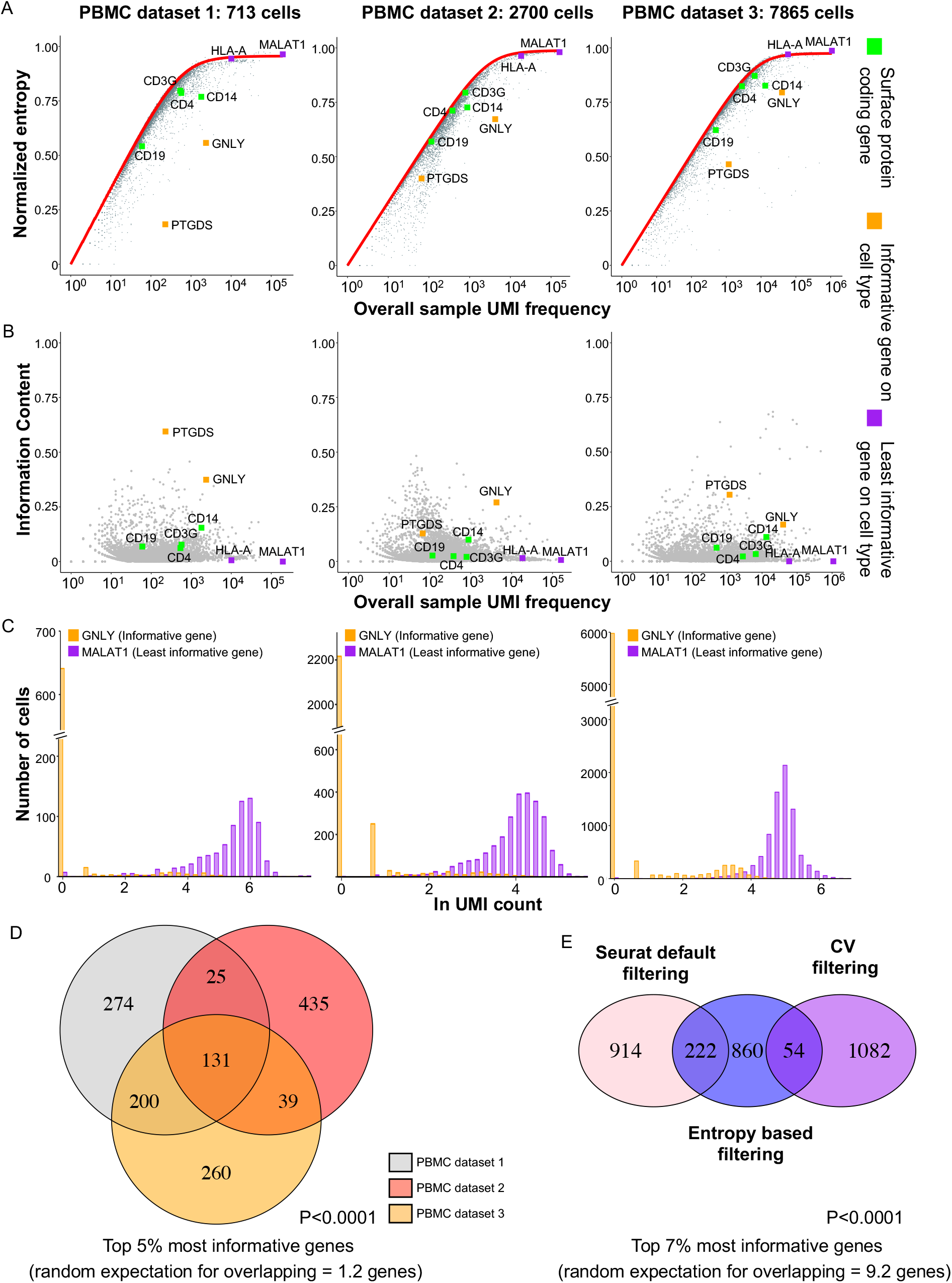
Context-specific identification of genes that are uninformative for cell type classification. We consider 3 PMBC datasets published by 10x Genomics. PBMC dataset 1 is the dataset we consider in the previously presented figures. PBMC dataset 2 and 3 comprise 2,700 and 7865 PBMCs from human blood samples (https://support.10xgenomics.com/single-cell-gene-expression/datasets/3.0.0/pbmc_10k_protein_v3 and https://support.10xgenomics.com/single-cell-gene-expression/datasets/1.1.0/pbmc3k). A. The conditional entropy across cells enables the identification of genes that cannot provide information for a classification task. The red lines show the expected entropy for genes that are randomly distributed across cells. The conditional entropies are normalized by *H*_*max*_. Most genes have distributions with entropies very close to the random expectation, implying that those genes cannot provide any information concerning cell classification. Two such uninformative genes are *MALAT1* and *HLA-A*, which code for highly-conserved proteins and are constitutively expressed in almost all cell types. In contrast, *GNLY*, which codes for a T-cell activation protein, and *PTGDS*, which codes for a protein involved in platelet aggregation, are quite informative about blood cell type. Most of the surface protein coding genes such as CD16 or CD19 are close to the null expectation and thus provides no additional information for cell type assignment. B. Distribution of information content for genes with different UMI frequency. Each dot represents a gene from PBMC dataset 1 (left), dataset 2 (middle), dataset 3 (right). The information content is calculated by the difference between randomly distributed null model and actual conditional entropy. C. Distribution of UMI counts for *GNLY* (orange) and *MALAT1* (purple) gene across cells in PBMC dataset 1 (left), dataset 2 (middle), dataset 3 (right). *MALAT1* is highly expressed in most cells and has a uniform distribution. *GNLY* is highly expressed in a small fraction of the cells and has a bimodal distribution. D. Overlap of the most informative genes from three datasets of peripheral blood mononuclear cells. The expected overlap across the three sets if we had simply selected 630 mRNAs at random from 12,600 would have been only about 1.2 mRNA on average, whereas overlap between two sets would have been above for most informative genes. Suggesting the robustness of the approach, and its biological significance, we find for both most and least informative mRNAs a statistically significant excess in the number of mRNAs that overlap across the three sets (p-values were calculated using non-parametric test with10,000 iterations). E. Overlap of most informative genes using Seurat default filtering approach, CV based filtering approach and our entropy-based approach using PBMC dataset 1. We select the same number of top informative genes from three filtering outputs.

A few genes such as *GNL*Y, which codes for a T-cell activation protein, and *PTGDS*, which codes for a protein involved in platelet aggregation, are informative about cell type across the three datasets (Fig. 3A, B). Importantly, we observe a large and consistent number of overlaps for the set of top 5%, 10% and 20 % most informative genes for cell type classification across the three datasets (Fig. 3D and Fig. S3B-D). We also observe a large overlap of informative genes using the default filtering algorithm in Seurat and our information-based filtering but not when we consider the coefficient of variation filtering approach (Fig. 3E). These findings suggest that the genes informative of cell type will be generally conserved across biological replicates. Importantly, because this approach requires no prior biological knowledge and contains no fitted parameters, it can <u>be applied without adjustment to samples from different tissues or from different organisms</u> (Fig. S3A).

We next compare the impact of using three feature filtering algorithms on the clustering accuracy of Seurat: coefficient of variation, default filtering algorithm in Seurat, and our information-based algorithm (Fig. 4A). Our analysis shows that using the coefficient of variation for filtering features performs poorly at detecting uninformative features. In contrast, the default Seurat filtering and our information-based filtering are truly able to select informative features. A point of distinction between the filtering algorithm implemented with Seurat and our proposed information-based filtering is their theoretical foundations. The filtering algorithm in Seurat algorithm models the mean-variance relationship within the data, as such is limited by the scope of its assumption and by the need to select parameter values. Information theory provides the best-grounded non-parametric approach to the problem of selecting informative features. Moreover, it requires the fewest assumptions and provides a roadmap for improvement and generalization. To our best knowledge, <u>it is unlikely that any *ad hoc* approach being proposed now and not grounded on information theory would be superior to that provided by information theory</u>.

**Figure 4.**
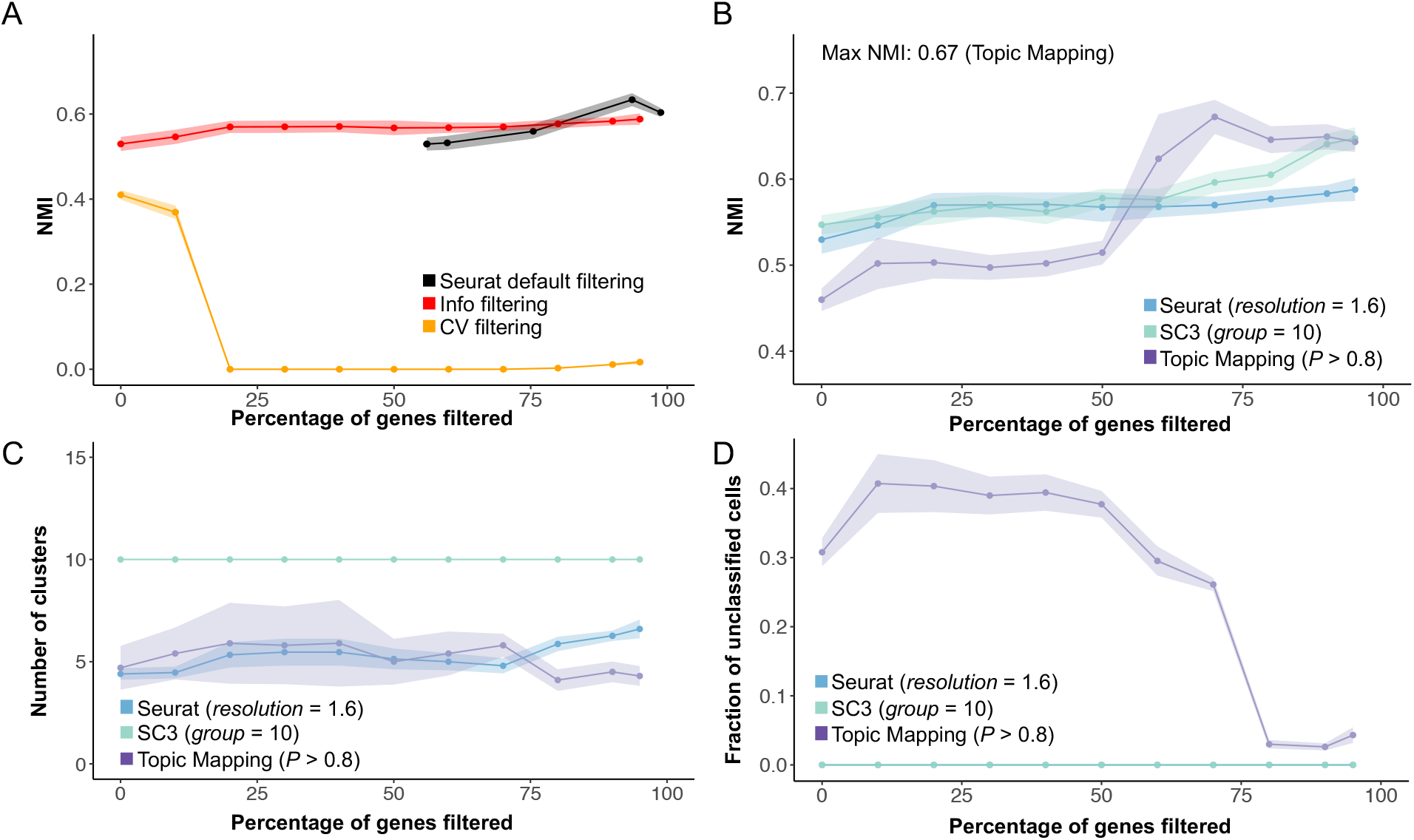
Filtering uninformative genes yields dramatic improvements in classification accuracy. A. Effect of filtering non-informative genes on accuracy using three different filtering approaches. We filter the genes using Seurat default filtering, our info filtering and coefficient of variance filtering and test the effects of classification accuracy on Seurat algorithm. B. Effect of filtering non-informative genes on accuracy of two standard algorithms (Seurat, SC3) and a new algorithm (Topic Mapping) for optimal parameter settings. We rank genes by the information content of the distribution and filter out an increasing percentage of the least informative genes. We generate 15 replicates using bootstrapping (sampling with replacement) in order to estimate confidence intervals for the accuracy of the different classification algorithms. We observe that there is no significant increase in accuracy until we remove at least 50% mRNAs, indicating that most mRNA measured in scRNA-seq screens contain no useful information for cell classification. As we then increase number of non-informative gene filtered out, we reveal a steady increase in accuracy across all testing algorithms regardless of parameter settings (see Fig. S4A). We also plot the results using Seurat default filtering algorithm. We also plot the results using Seurat default filtering algorithm. C. Filtering out non-informative genes does not alter the estimated number of detected clusters. D. Filtering out non-informative genes reduces the percentage of unclassified cells in Topic Mapping. The number of unclassified cells reduce significantly after we remove 50% of mRNAs for Topic Mapping. For this dataset there are no unclassified cells in SC3. Seurat always classifies every cell.

We next evaluate the impact of using our filtering algorithm on clustering accuracy of three algorithms: SC3 (Kiselev et al. 2017), Seurat (Satija et al. 2015), and Topic Mapping (Lancichinetti et al. 2015). Topic Mapping is a highly reproducible high-accuracy algorithm for clustering documents. We implement a straightforward analogy: words to genes and documents to cells.

As the fraction of filtered genes increases, the classification accuracy of all three algorithms also increases (Fig. 4B and Fig. S4A). Interestingly, the greatest gain in accuracy occurs once we remove the 50% genes with the lowest *I*(*g*), demonstrating that most genes measured in scRNA-seq truly contain no useful information for cell classification. Indeed, clustering accuracy increases up to the removal of the bottom 90% of genes by information content.

Filtering out uninformative genes greatly improves classification stability (Fig. 4C-D and Fig. S4B). Specifically, the number of clusters returned by Seurat and Topic Mapping become more stable as the percentage of genes filtered increases. Additionally, the confidence with which cluster assignment are made in Topic Mapping also increase with percentage of filtered genes. Importantly, the genes that are most informative in our analysis are dissimilar to those used for cellular classification based on protein measurements (Fig. 3A). This suggests that classification schema based on genes coding for proteins targeted in flow cytometry, immunohistochemistry or other protein-based measures (Bhattacharya et al. 2014), will not necessarily extract the optimal amount of information from scRNA-seq data.

Our results also suggest that “noise” from uninformative genes leads to ambiguous assignment of cells to clusters. After removing uninformative genes, the three algorithms largely correctly assign major cell groups such as B cells or monocytes (Figs. 5A, C, E). Seurat and SC3, but not by Topic Mapping, also correctly identify NK cells. However, all three have trouble distinguishing between other cell types. CD4, CD8 and NKT cells are placed into the same clusters by the three methods, but while SC3 and Seurat arbitrarily break the cells from those types into two clusters, Topic Mapping puts them all in a single cluster. The confusion matrices are similar when we compare the same clustering results against a supervised annotation approach using SingleR and ENCDOE or HPCA as reference dataset (Figs. 5 B, D, F and Fig. S5C). While the supervised annotation approach identifies boarder cell groups, our protein-based annotation significantly overlaps with their results (Figs. S5A, B).

**Figure 5.**
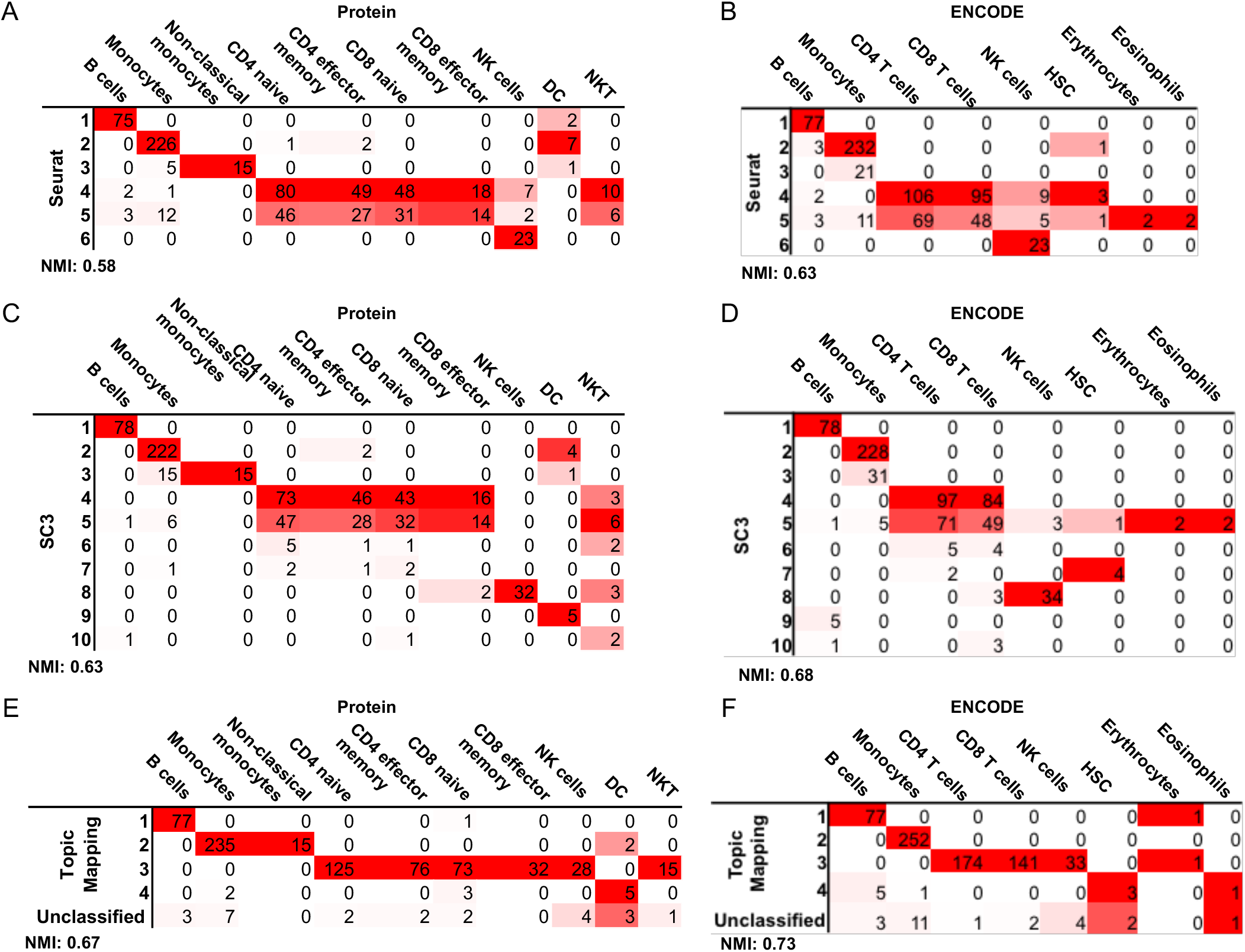
Confusion matrices for Seurat, SC3 and Topic Mapping classification of mRNA with 80% filtering against surface-protein based classification (A, C, E) and ENCODE based annotation (B, D, F), respectively. For Topic Mapping, unclassified cells are listed in a separate row and excluded from NMI calculation.

### Study Limitations

There are limitations in our study. Our benchmarking method is limited to CITE-seq platform where both surface protein and mRNA are quantified at the same time. The number of CITE-seq datasets is limited in current literature. However, CITE-seq analysis is becoming increasing popular in medicine (Saigusa and Ley 2020; Bandyopadhyay et al. 2019; Kotliarov et al. 2020) and our method can be readily applied to these new datasets. On the other side, our filtering approach is not limited to CITE-seq platform and can be applied in any scRNA-seq data without any prior biology knowledge. Secondly, we recognize that researchers perform some filtering procedures to remove the undetected genes before clustering algorithms as best practice. However, most of current filtering rely on parametric tests, there could be potential biases in the assumptions. Our method, on the other hand, does not reply on parametric test and has minimal assumptions. Lastly, there exist supervised cell annotation tools such as SingleR (Aran et al. 2019). We demonstrate our annotation method largely overlaps with the results based on ENCODE and HPCA reference but with more details. Furthermore, our labels can be used as a reference dataset in those supervised approach.

## Discussion

Data sparsity due to low RNA capture rates and uninformative genes are frequently acknowledged as challenges in the analysis of scRNA-seq data (Kharchenko, Silberstein, and Scadden 2014). Depending on the sequencing platform, experimental setup, and sample preparation, on the order of 80% of the scRNA-seq measurements for each of the approximately 20,000 human genes yield zero counts (Angerer et al. 2017). Confirming prior research, we find that high sparsity and the large number of uninformative genes significantly reduces the performance of clustering algorithms. We show that this reduction in clustering accuracy and reproducibility can be addressed using a quantitative approach based on information theory. We show this approach provides a quantitative assessment of the information content for cell classification of any gene in a specific dataset without the need for any prior biological knowledge. When applied to a benchmarking data set generated independently of RNA-seq data, this method provides a significant improvement in classification accuracy when compared to other widely used tools (Kiselev, Andrews, and Hemberg 2019).

Our study highlights the importance of independent validation for benchmarking scRNA-seq analysis tools to avoid circularity and lack of reproducibility. The entanglement of cell type assignment with assessment of clustering quality carries an inherent risk of confirmation bias. This situation is exacerbated by the widespread practice of using visual inspection to confirm clustering quality, in which the authors use the specificity of gene assignment to a given cluster (feature plots) to support the accuracy of the clustering algorithm. Information theory based approaches offer an objective alternative by providing a quantitative measure of the information conferred by each gene for determining cellular annotations. We are hopeful this approach will be applied by others to additional benchmark datasets with a goal of increasing reproducibility and accuracy of scRNA-seq analysis.

## Methods

### Benchmarking CITE-seq protein data with decision tree modeling

We model the distribution using a Gaussian mixture model with two peaks and determine a binary cutoff by identifying the value for which the likelihoods of a cell belonging to each of the two groups are identical. We list the estimated parameters for the decision tree in the supplementary information.

In order to test the robustness of our classification and to acknowledge uncertainty in a cell’s classification, we also consider the case in which we set cells with protein levels that yield odds ratios smaller than 5:1 between estimated Gaussian functions for two peaks as “unclassified cells”.

### Clustering PBMC scRNA-seq data

#### Seurat

We download version 3.2.1 of the Seurat package from CRAN R project and use default defaults settings illustrated by the online tutorial (https://satijalab.org/seurat/vignettes.html). Different parameters in resolution are sampled to reflect the change in clustering accuracy.

#### SC3

We download version 1.16.0 of the SC3 package from Bioconductor project and use default settings except for the selection of the number of clusters. Several cluster numbers are used to examine the clustering accuracy.

#### Topic Mapping

We obtained the Topic Mapping package from the authors. We implement a straightforward analogy: words to genes and documents to cells. The TopicMapping algorithm outputs two conditional probability vectors: p(gene|topic), which is used to identify marker genes for each cluster, and p(topic|cell), which is used to identify cell clusters based on probability distribution over topics. Each topic is treated as a cell type and the cells are assigned to the topic with the highest probability. There are only two parameters in determining the clustering results *P*, which specifies the statistical significance threshold for acceptance of new clusters, and *t*, which specifies which required minimal manipulation. We use default values (*P* = 0.05 and *t* = 10) in our analyses.

### Generating synthetic datasets to model sparsity and uninformative genes

We model data sparsity by setting a model for the probability that a specific surface protein’s level is too low to be detected. We use a sigmoidal function to generate the probability that protein *j* is detected by the assay for an individual cell *c*:

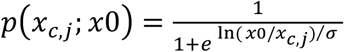

where *x* is the actual protein level, ln is the natural logarithm function, *x*0 is the selected detectability threshold, and *σ* controls the stochasticity of the detection process around the threshold. For simplicity, we set *σ* = 1 for all simulations.

To model the impact of uninformative genes on clustering accuracy, we add the data from 500, 1000, 2000, 8000, 16,000 randomly selected genes from the transcriptomics dataset to the synthetic data for protein levels with distinct detectability threshold generated in the previous step.

### Calculating the information content of all genes for a sample

Our method is generalized from that reported by (Gerlach, Shi, and Amaral 2019). Consider a sample *S* comprised of *L* cells and denote by *n*(*g*, *c*) the number of UMIs for gene *g* and cell *c*. The total reading depth *N* can be expressed as the sum over all genes *g* and all cells *c* of *n*(*g*, *c*). The probability that a UMI for gene *g* occurs in cell *c ∊ S* is:

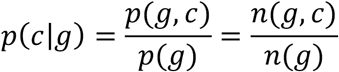

Where *p*(*g*) = *N*(*g*)/*N* is the relative frequency of UMIs for gene *g* across the entire sample. We can use Shannon entropy to quantify the heterogeneity of this distribution:

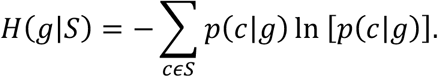

The entropy *H*(*g*|*S*) thus describes the information we can hope to extract from the distribution of number of UMIs for gene *g*. A gene specific to a certain cell type will have less uncertainty in its expression pattern across the sample and thus will have a lower entropy than a ‘house-keeping’ gene which will not be specific to a certain cell type. The maximum possible value of the conditional entropy occurs for a uniform distribution of number of UMIs,

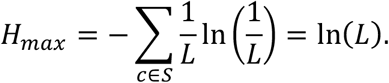

Due to high sparsity nature of scRNA-seq dataset, most genes will occur at very low frequencies and the limit expressed by *H*_*max*_ will not be accurate. Therefore, we estimate a null model for the *n*(*g*) in order to calculate the expected value of the entropy under the null hypothesis. To this end, we generate random distributions of UMI counts across genes and cells while preserving the marginal counts *N*(*g*) and *N*(*c*) = ∑_g_ *n*(*g*, *c*). By calculating the expected entropy from many realizations of this null model, we define the “true” information content of a gene, *I*(*g*), as:

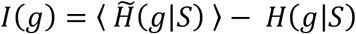

where 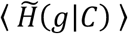 denotes the average conditional entropy over different realizations of the null model, and higher *I*(*g*) indicates higher information-content. In contrast, genes with *I*(*g*) ≈ 0 are uninformative and should be excluded from the list of features during the clustering analysis.

### Comparison of protein-based cell type annotation with supervised methods

We compare our protein-based cell type annotation with the supervised cell type annotation method using SingleR with both Human Primary Cell Atlas (HPCA) and Encyclopedia of DNA Elements (ENCODE) dataset. SingleR package is obtained through CRAN R project and default parameters are used in cell type annotation for single cells.

## Data and code availability

We use a publicly available peripheral blood mononuclear cells (PBMC) dataset compiled by 10x Genomics: PBMC dataset 1 (https://support.10xgenomics.com/single-cell-gene-expression/datasets/3.0.0/pbmc_1k_protein_v3). PBMC dataset 2 (https://support.10xgenomics.com/single-cell-gene-expression/datasets/3.0.0/pbmc_10k_protein_v3) and PBMC dataset 3 https://support.10xgenomics.com/single-cell-gene-expression/datasets/1.1.0/pbmc3k). Human and mouse pancreas cells data were downloaded on 04/03/2018 from https://hemberg-lab.github.io/scRNA.seq.datasets/human/pancreas/.

The source code is publicly available at https://github.com/amarallab/Benchmark_scRNA_seq.

## Statistical analysis

All statistical analyses are performed using R (Team 2014) and figures are produced using the package ggplot2 (Wickham). The error bar and confident intervals are calculated using bootstrapping. P-value is calculated using student’s t-test with two side for figure 2 and hypergeometric test for Venn diagram in figure 3 and figure S3 using build-in function in R.

## Acknowledgements

Z.R. is supported from American Heart Association Postdoctoral Fellowship 20POST35180141. LANA is supported by the Successful Clinical Response In Pneumonia Therapy (SCRIPT) Systems Biology Center (U19AI135964); and the Northwestern University Quantitative Biology Center (NUQuB) (NSF/Simons DMS-1764421);

---

**Figure S1.**
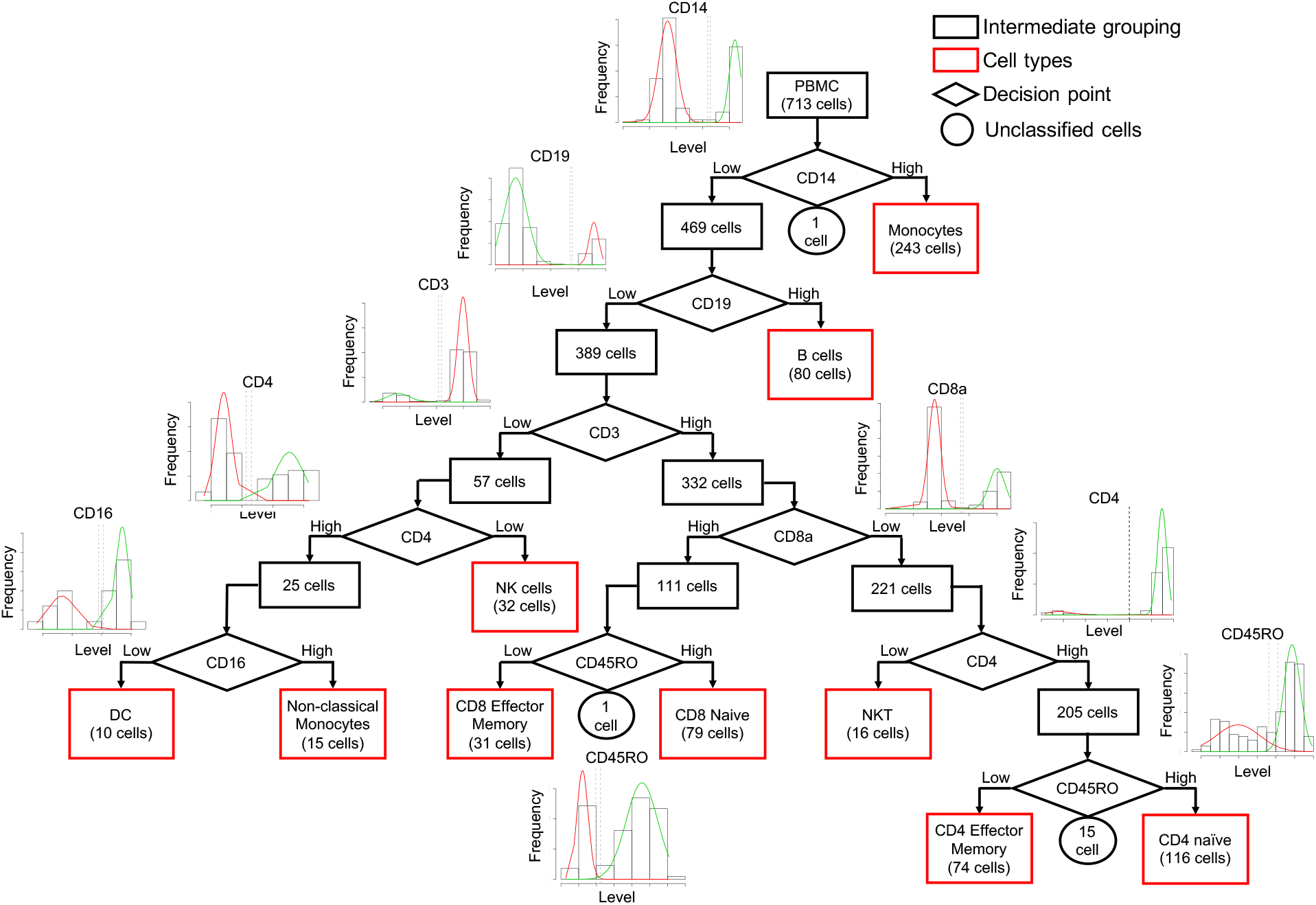
Creation of externally validated labelling for peripheral blood mononuclear cells (PBMCs) with uncertainty thresholds. We model the empirical distribution as a mixture of two Gaussian peaks, which can be used for classification with a biologically-grounded decision tree enabling us to classify every cell into one of ten cell types.

**Supplementary Table 1. The estimated parameters for each decision tree using Gaussian mixture model.** (Data shown in separated file)

**Supplementary Table 2. Cell type classification based on binary cutoffs.** (Data shown in separated file)

**Supplementary Table 3. Cell type classification based on uncertainty regions.** (Data shown in separated file)

**Figure S2.**
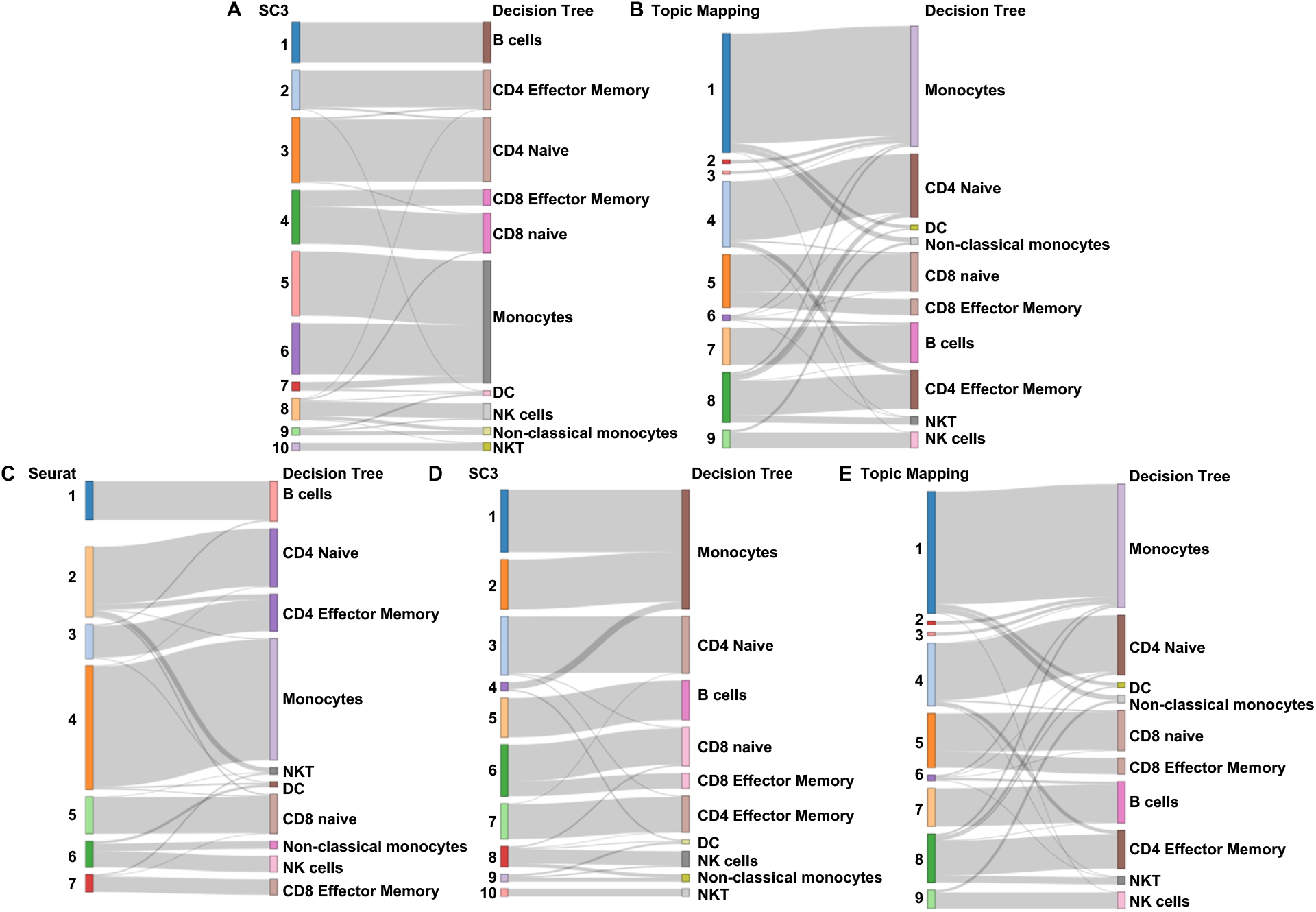
Sankey diagram of classification based on surface protein levels using uncertainty region. For the first two panels, the right side shows the cell types we identify using binary cutoffs of surface proteins. The left side shows the clusters identified by A. SC3 algorithm with the user input of correct cluster number (10); B. Topic Mapping algorithm. For the last three panels, the right side shows the cell types we identify using binary cutoffs of surface proteins. The left side shows the clusters identified by C. Seurat algorithm at default *resolution* (0.8); D. SC3 algorithm with the user input of correct cluster number (10); Topic Mapping algorithm.

**Figure S3.**
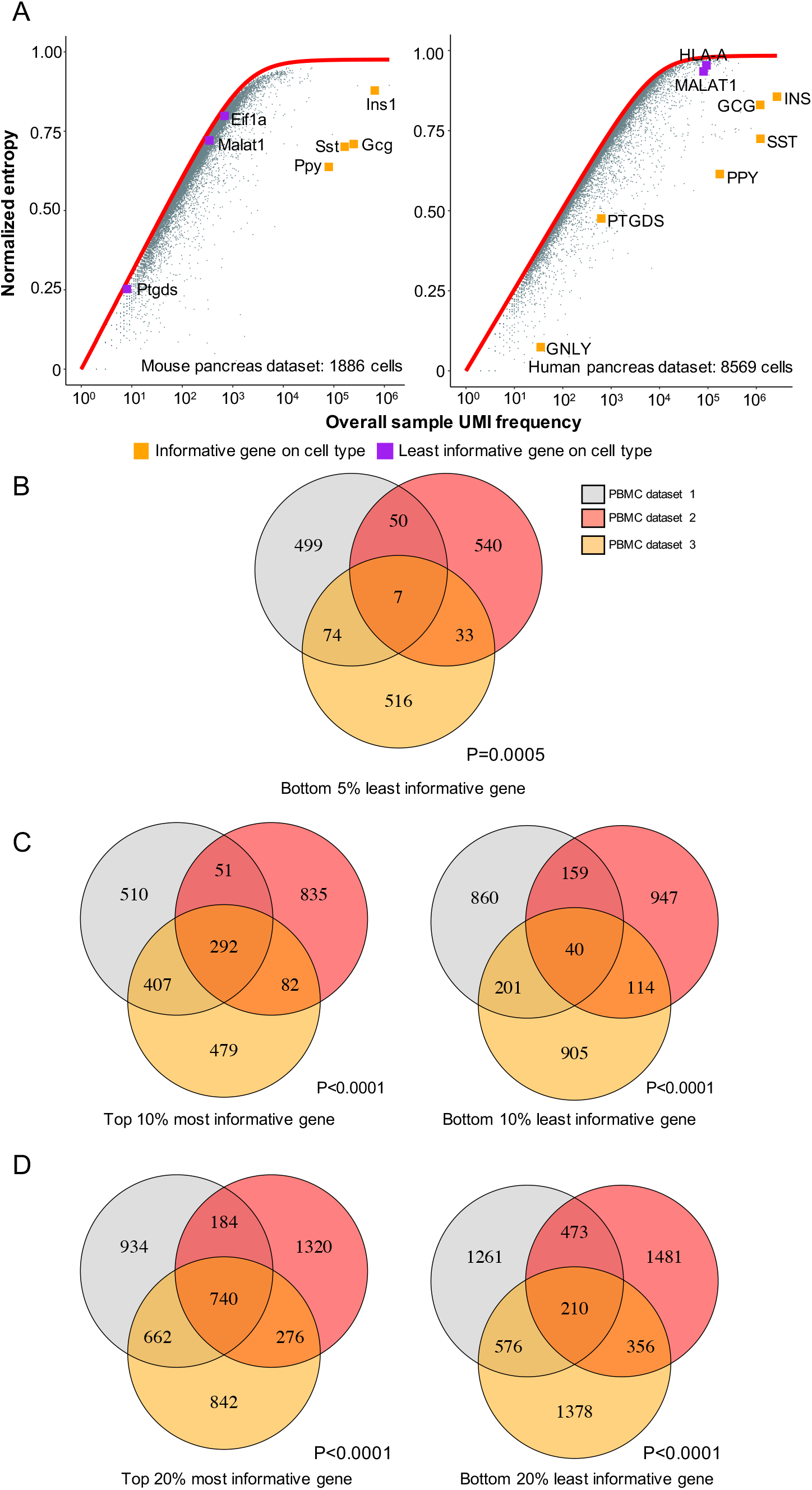
Context-specific identification of genes that are uninformative for cell type classification. A. The conditional entropy across cells does not require *a priori* biological knowledge in determining uninformative genes. We consider a published dataset of pancreas cells from either human or mouse and calculate the information content for each gene. Genes such as INS that are critical for pancreas cell types identification are with high information content. *MALAT1* or *HLA-A* are uninformative in both human and mouse datasets like what we find in PBMC datasets. Some genes such as *PTGDS* are informative for human cell types but not for mouse. B-D: Overlap of most informative and least informative genes (B: 5%; C:10%; D: 20%) using information contents from three datasets of peripheral blood mononuclear cells.

**Figure S4.**
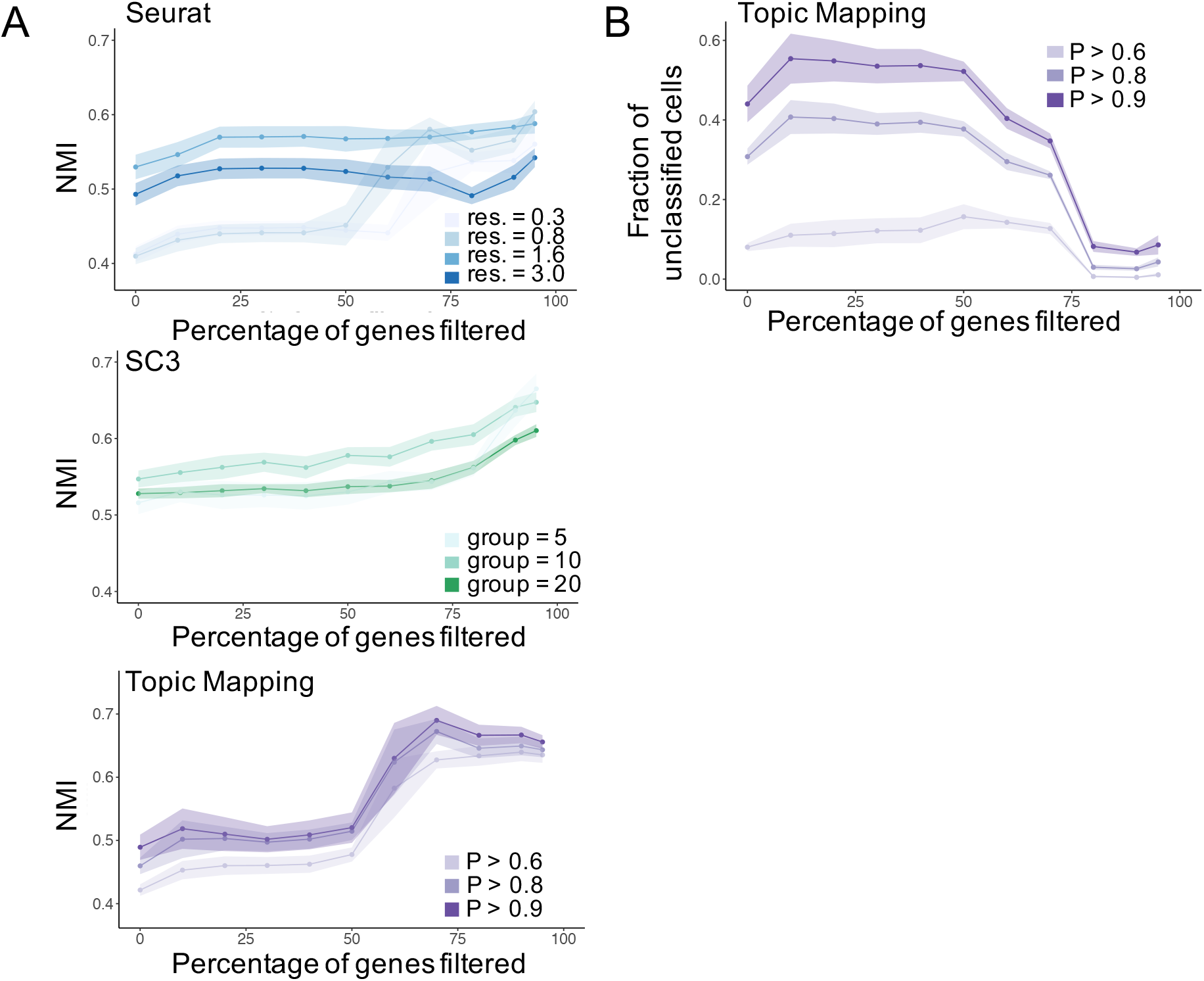
Filtering uninformative genes yields dramatic improvements in classification accuracy. A. Effect of filtering non-informative genes on accuracy of two standard (Seurat, SC3) and a new (Topic Mapping) algorithms for different parameter settings. B. Filtering out non-informative genes reduces the percentage of unclassified cells in Topic Mapping for different parameter settings.

**Figure S5.**
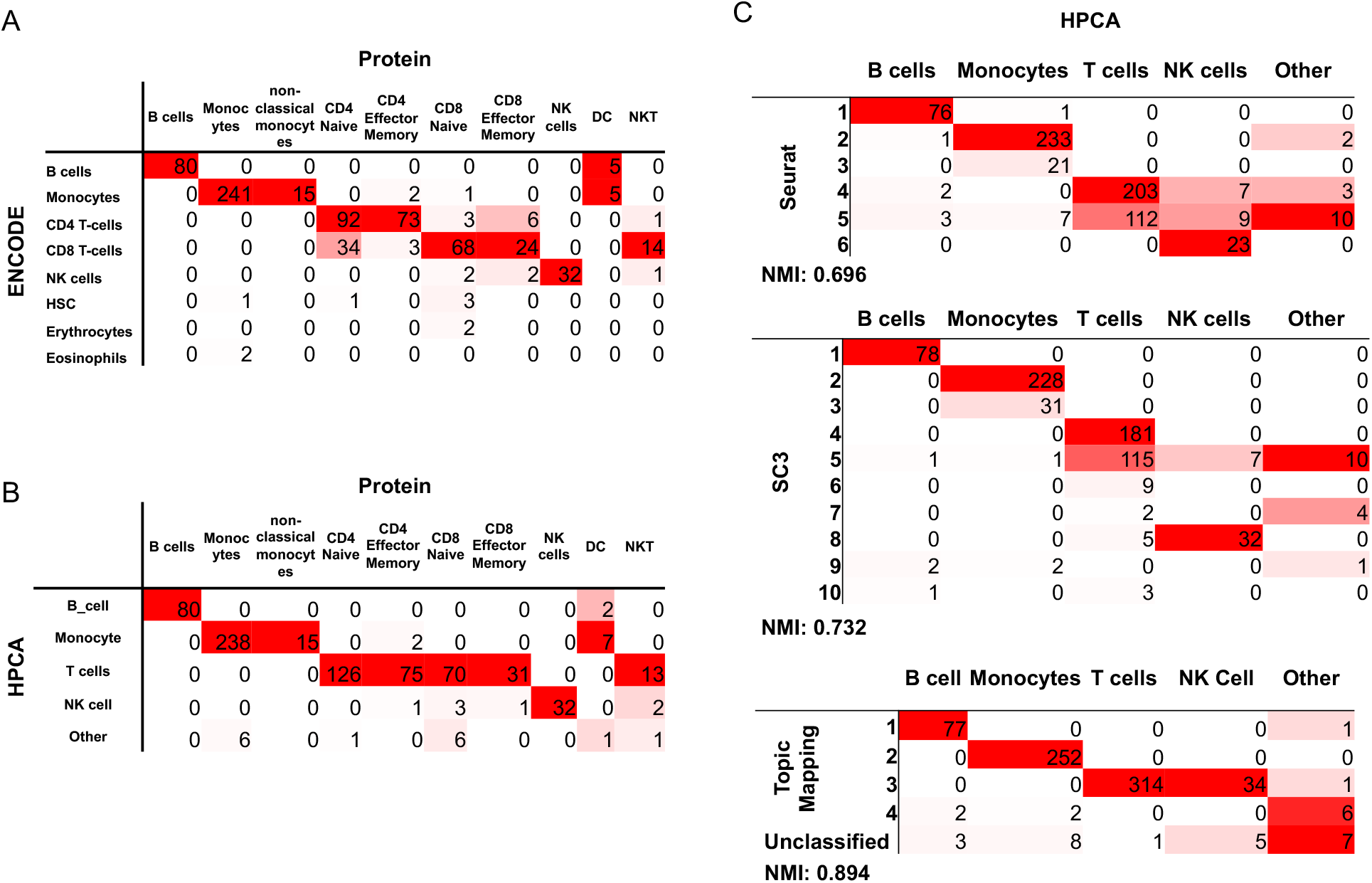
Supervised annotation methods highly overlap with our protein-based annotation. A. Confusion matrix for supervised annotation method using ENCODE SingleR and our protein-based annotation. B. Confusion matrix for supervised annotation method using HPCA SingleR and our protein-based annotation. C. Confusion matrix for Seurat (top), SC3 (middle) and Topic Mapping (bottom) classification of mRNA with 80% filtering against HPCA based annotation.

